# The herpes simplex virus 1 protein ICP4 acts as both an activator and repressor of host genome transcription during infection

**DOI:** 10.1101/2021.04.09.439230

**Authors:** Thomas Rivas, James A. Goodrich, Jennifer F. Kugel

## Abstract

Infection by Herpes simplex virus 1 (HSV-1) impacts nearly all steps of gene expression in the host cell. The regulatory mechanisms by which this occurs, and the interplay between host and viral factors, have yet to be fully elucidated. Here we investigated how the occupancy of RNA polymerase II (Pol II) on the host genome changes during HSV-1 infection and is impacted by the viral immediate early protein ICP4. Pol II ChIP-seq experiments revealed a reduction of Pol II occupancy across the bodies of hundreds of host genes that was dependent upon ICP4. Concomitantly, Pol II levels increased across the bodies of several hundred genes, the majority of which also depended on ICP4 for activation. Our data suggest ICP4 regulates repression of Pol II at host genes by inhibiting recruitment of Pol II, while it regulates activation by promoting release of Pol II from promoter proximal pausing into productive elongation. Consistent with this, relative levels of the pausing factors NELF-A and Spt5 were reduced on an HSV-1 activated gene in an ICP4 dependent manner. Exogenous expression of ICP4 revealed that ICP4 can activate, but not repress, transcription of some genes in the absence of infection in a manner that correlates with the chromatin state of the gene. Together our data support the model that ICP4 decreases promoter proximal pausing on host genes activated by infection, and ICP4 is necessary, but not sufficient, to repress transcription from host genes during viral infection.

## Introduction

Herpes simplex virus 1 (HSV-1) is a large double-stranded DNA virus that encodes approximately 80 proteins expressed in a temporally regulated cascade upon infection of a host cell, categorized as the immediate early (IE) proteins, early proteins, and late proteins [1]. There are five IE proteins (ICP0, ICP4, ICP22, ICP27, ICP47) expressed rapidly after infection, and all but ICP47 act to prevent the host cell from silencing the viral genome and to facilitate transcription of the early and late viral genes [2–5]. The early gene products largely promote viral DNA replication, and the late gene products are necessary for creating viral progeny and include the structural components of the virus [6, 7]. None of the ∼80 viral genes code for an RNA polymerase; therefore, HSV-1 hijacks the host cell RNA polymerase II (Pol II) and general transcription factors to transcribe its own genome.

Coincident with the expression of viral genes is a substantial reduction in the levels of host mRNA transcripts, which results from both decreased transcription of host genes as well as mRNA degradation by the viral host shutoff protein, vhs [8–11]. Consistent with decreased levels of host genome transcription, studies have shown there is a widespread reduction of Pol II occupancy on the host genome within hours after HSV-1 infection [12]. Studies have also implicated several IE viral proteins in influencing or associating with Pol II or its general transcription factors [13–17], although the impact on host transcriptional activity is not clear. A recent study identified the IE protein infected cell polypeptide 4 (ICP4) as a major regulator of Pol II occupancy at host promoters facilitating the transcriptional switch from the host genome to the viral genome [18]. Recent studies have also shown that ICP27 is important for regulating transcription termination by Pol II at host genes [13, 17]. Much remains to be learned about the interplay between host and viral factors in regulation of Pol II activity during infection.

mRNA transcription in human cells is a highly regulated process that requires the coordinated activity of many protein factors. Transcription requires chromatin remodelers and histone modifiers for Pol II to navigate through chromatin [19]. Pol II is first recruited to the promoters of genes with the help of general transcription factors TFIIA, TFIIB, TFIID, TFIIE, TFIIF, TFIIH, and the Mediator complex to form pre-initiation complexes (PICs) [20]. Gene specific activator and repressor proteins that bind at enhancers help control the assembly of PICs on core promoters [21]. After transcription initiates Pol II pauses 20-100 nucleotides downstream of the start site, a step known as promoter proximal pausing, where Pol II can either undergo premature termination or proceed into the gene body to productive transcript elongation [22]. Promoter proximal pausing is controlled by the activities of the pausing factors DSIF (DRB sensitivity inducing factor) and NELF (negative elongation factor), which associate with Pol II at the promoter proximal pause [23]. The kinase subunit of PTEF-b (positive transcription elongation factor) phosphorylates NELF and DSIF, inducing conformational changes to release NELF from the paused polymerase and facilitates the transition of Pol II into an elongation-competent conformation [23].

Also important for early steps in Pol II transcription are post-translational modifications on the C-terminal domain (CTD) of the largest Pol II subunit, which is comprised of 52 heptad-repeats with the consensus sequence YSPTSPS. The CTD is differentially phosphorylated during transcription, with Ser5 phosphorylation (Ser5-P) enriched around the transcriptional start site (TSS) and splice sites, and Ser2 phosphorylation enriched throughout gene bodies and surrounding the poly-adenylation site at the 3’ ends of genes [24]. Transcription of the viral genome by Pol II occurs through the same regulatory steps as transcription of the host genome: PIC assembly, promoter proximal pausing, elongation, then termination, although an interplay of viral proteins and host cell transcription machinery are involved [13,25–27]. In particular, ICP4 is critical for viral transcription. It is found coating the viral genome early during infection and is thought to facilitate the recruitment of host machinery [18].

ICP4 is an essential viral protein that is required for the transcription of early and late viral genes [28, 29], and in the absence of ICP4 viral DNA replication is severely impaired [30]. ICP4 binds to the HSV-1 genome with a loose consensus sequence, however, has also been shown to bind promiscuously [18, 31]. ICP4 interacts with the host transcription factors TFIID and Mediator and is thought to help recruit both factors to the viral genome [32]. Indeed, data suggest that ICP4 strengthens the interaction of TFIID with viral promoter DNA to facilitate PIC formation and more robust initiation of viral transcription [25]. The role of ICP4 as a transcriptional regulator has largely been studied in the context of viral transcription. Recently, however, ICP4 ChIP-seq data showed that after HSV-1 infection, ICP4 is also present on the host genome near transcriptional start sites [18]. There was little correlation between sites of ICP4 occupancy and the consensus sequence that ICP4 binds, suggesting a non-sequence specific presence on the host genome. Furthermore, lytic infection by HSV-1 caused a decrease in occupancy of Ser5-P Pol II at TSSs of host cell genes, and infection with a mutant virus strain that codes for a truncated, nonfunctional form of ICP4 (HSV-1 n12 strain) showed recovery of Ser5-P Pol II occupancy at TSSs [18]. These data suggest that ICP4 is required for the loss in Ser5-P Pol II on host cell genes during infection. The mechanisms by which this occurs, and how ICP4 impacts host cell transcription at activated versus repressed genes remain to be investigated.

We are interested in understanding mechanisms by which host cell transcription is controlled during HSV-1 infection, including the functions of ICP4. Toward these goals, we evaluated the effects of infection by wild type (WT) HSV-1 and n12 HSV-1 (ICP4 defective) on total Pol II occupancy across the human genome using ChIP-seq. For infection by each virus, we identified genes whose Pol II levels were increased (activated) or decreased (repressed) compared to mock infection. WT infection activated and repressed Pol II levels across the bodies of hundreds of host genes. n12 infection resulted in little repression of Pol II levels; however, activated Pol II levels at a similar number of host genes compared to WT infection, albeit to a lesser extent. Further investigation of the Pol II ChIP-seq data revealed that ICP4 promotes release of promoter proximally paused Pol II at genes activated during infection, while negatively impacting Pol II recruitment to promoters of genes repressed during infection.

Expression of ICP4 in the absence of infection activated transcription of some host genes as well as transiently transfected luciferase reporter plasmids, which was decreased in stable cell lines containing the reporter plasmids. We propose that ICP4-mediated activation is sensitive to the chromatin state of the gene. Together, our data support dual activities for ICP4 in regulating host genome transcription during HSV-1 infection: broadly reducing Pol II occupancy on repressed genes and altering promoter proximal pausing at activated genes.

## Results

### The HSV-1 protein ICP4 is required for host cell transcriptional reprogramming upon HSV-1 infection

With the goal of understanding how host cell transcription is regulated during HSV-1 infection, and the role of ICP4, we determined changes in Pol II occupancy on the host genome during viral infection using Pol II ChIP-seq. Human HEK 293 cells were infected for four hours at a MOI of 10 with WT virus (HSV-1 KOS strain), or n12 virus (a mutant HSV-1 virus that codes for a truncated, nonfunctional ICP4) [33]. Infected and mock infected cells were formaldehyde crosslinked and the chromatin was isolated and sheared by sonication. The sheared chromatin was used in ChIP assays with an antibody against the N-terminal region of the largest subunit of Pol II (Rpb1), allowing for immunoprecipitation independent of the phosphorylation state of the Pol II CTD. Illumina sequencing libraries were prepared using the ChIP eluates from biological replicates, as well as input chromatin. The high-throughput sequencing reads were mapped to the human genome (other studies have analyzed changes in Pol II occupancy on the HSV-1 genome and the role of ICP4 [18,26,34]; hence here we focused on the host genome). The sequencing reads from biological replicates were tightly correlated, as shown by the correlation matrix and PCA plots in Supplemental Figure 1, therefore the sequencing reads from replicates were combined for much of the analyses.

Pol II occupancy on the human genome under the three conditions (i.e., mock, WT, n12) was first analyzed across the bodies of mRNA genes. For each gene, we compared the mapped sequence reads per kilobase (RPK), normalized to a constant sequencing depth, from +250 relative to the annotated TSS through the annotated 3’-end of the gene. By doing so we removed promoter proximally paused Pol II from consideration because changes in paused Pol II do not accurately reflect changes in the level of transcription of genes [23]. In addition, we only considered genes with average gene body Pol II occupancy levels at least 2-fold greater than the average gene body RPK in input chromatin. For the 8732 genes that fit these criteria across all three conditions, we generated box plots that show the quartiles of Pol II occupancy levels for each condition (Figure 1a). Infection with the WT virus triggered a statistically significant decrease in Pol II levels across the bodies of mRNA genes (compared to mock-infected cells), which agrees with prior studies [10, 12]. Infection with the n12 virus resulted in gene body Pol II levels that were not statistically different from the mock condition but were increased relative to the WT-infected cells. Therefore, the loss of total Pol II across the bodies of mRNA genes due to HSV-1 infection requires ICP4. This is consistent with prior results showing that loss of Ser5-P Pol II in the promoter proximal region of genes requires ICP4 [18].

**Figure 1.**
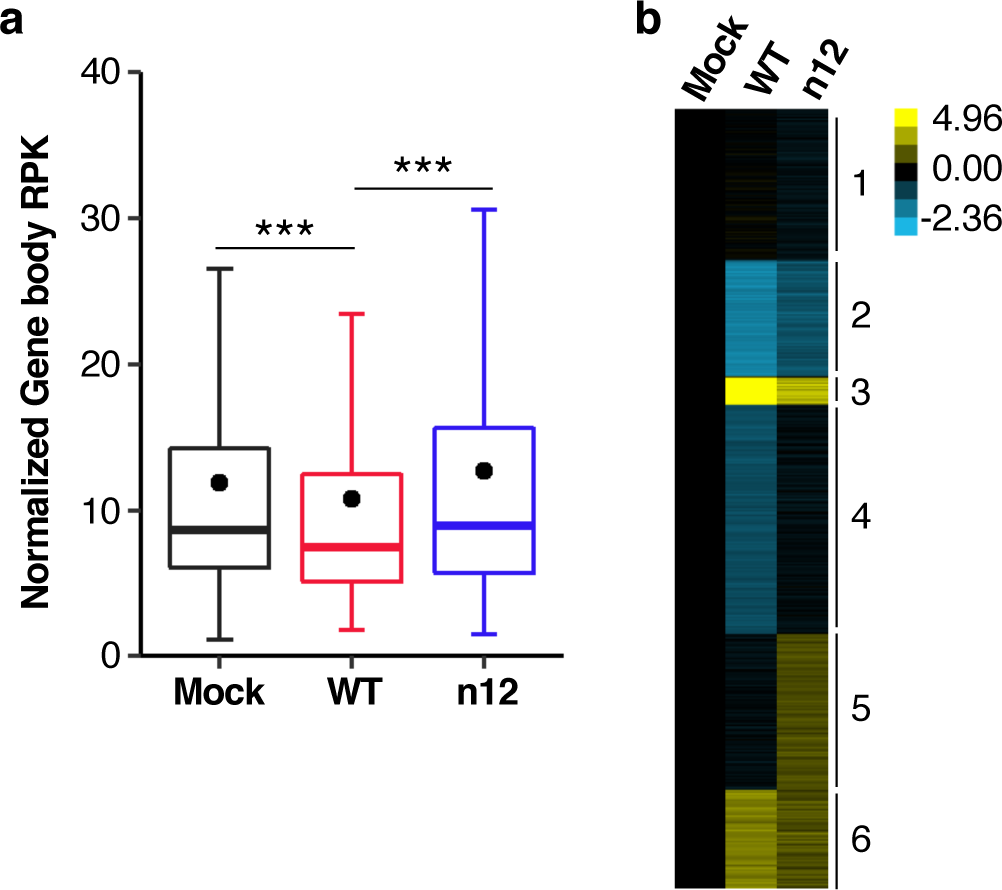
HSV-1 infection causes a global reprogramming of Pol II occupancy on the bodies of host cell mRNA genes, which is dampened in the absence of ICP4. **(a)** Pol II levels on cellular mRNA genes are largely reduced during HSV-1 infection in an ICP4 dependent manner. Plotted are the quartiles of Pol II ChIP-seq reads per kb in the gene bodies (+250 to the 3’ end, normalized to constant depth of sequencing) of RefSeq genes greater than 1kb and with Pol II levels 2-fold above the average input chromatin (8732 genes). The black dot represents the mean. P values were determined by an unpaired Mann-Whitney test with *** representing a p-value < 2.2x10^-16^. **(b)** Both increases and decreases in Pol II levels are observed at host cell gene bodies after infection with WT and n12 virus. The heat maps show the fold-change in gene body Pol II ChIP-seq reads compared to the mock condition for 8732 genes. Gene body RPK for WT and n12 infected data were normalized to the mock condition for each gene, log_2_-transformed, and clustered (k-Means, 6 clusters). Cyan represents a decrease and yellow represents an increase.

We next clustered genes according to the fold-change in their gene body Pol II levels after infection and plotted them as a heatmap in Figure 1b with each horizontal line representing an individual gene. In WT-infected cells, the Pol II changes were dominated by decreases (e.g., clusters 2 and 4) with a smaller subset of genes that increased upon infection (clusters 3 and 6). These changes were generally dampened during infection in the absence of ICP4; however, there was a group of genes whose Pol II levels specifically increased upon n12 infection (cluster 5). These clusters show that both increases and decreases in Pol II levels on the bodies of host cell mRNA genes are regulated in part by ICP4.

### ICP4 is required for the reduction of Pol II on host mRNA genes upon HSV-1 infection as well as the increase in Pol II at hundreds of other genes

We applied differential gene expression algorithms to identify mRNA genes that showed a statistically significant increase (i.e. activation) or decrease (i.e. repression) in Pol II gene body occupancy upon infection with either virus compared to the mock condition using DESeq2. Figure 2a shows MA plots for WT versus mock (top) and n12 versus mock (bottom). Genes were considered differentially expressed (black dots) if they had an adjusted p-value < 0.05 and a log_2_ fold change ≥ 1 (activated) or ≤ -1 (repressed). There were 463 and 402 activated genes for WT and n12 infection, respectively, and 626 and 57 repressed genes for WT and n12 infection, respectively. None of the genes repressed in WT infection were considered statistically significantly activated in n12 infection. The genes activated upon infection with either virus include genes that were transcribed in mock infected cells, as well as genes that did not have Pol II occupancy above input chromatin prior to infection, therefore, HSV-1 infection activates genes regardless of prior transcriptional activity. The overlap between the identities of the activated and repressed genes in the WT and n12 infections are shown as Euler diagrams in Figure 2b. This shows there are 294 host cell genes that rely on the presence of ICP4 for activation and 233 host cell genes that are uniquely activated by infection in the absence of ICP4. Similarly, there are 588 genes that rely on the presence of ICP4 for repression in WT infection and 19 genes that are uniquely repressed in the absence of ICP4. Deletion of ICP4 greatly reduced the number of repressed genes upon infection and altered the set of activated genes.

**Figure 2.**
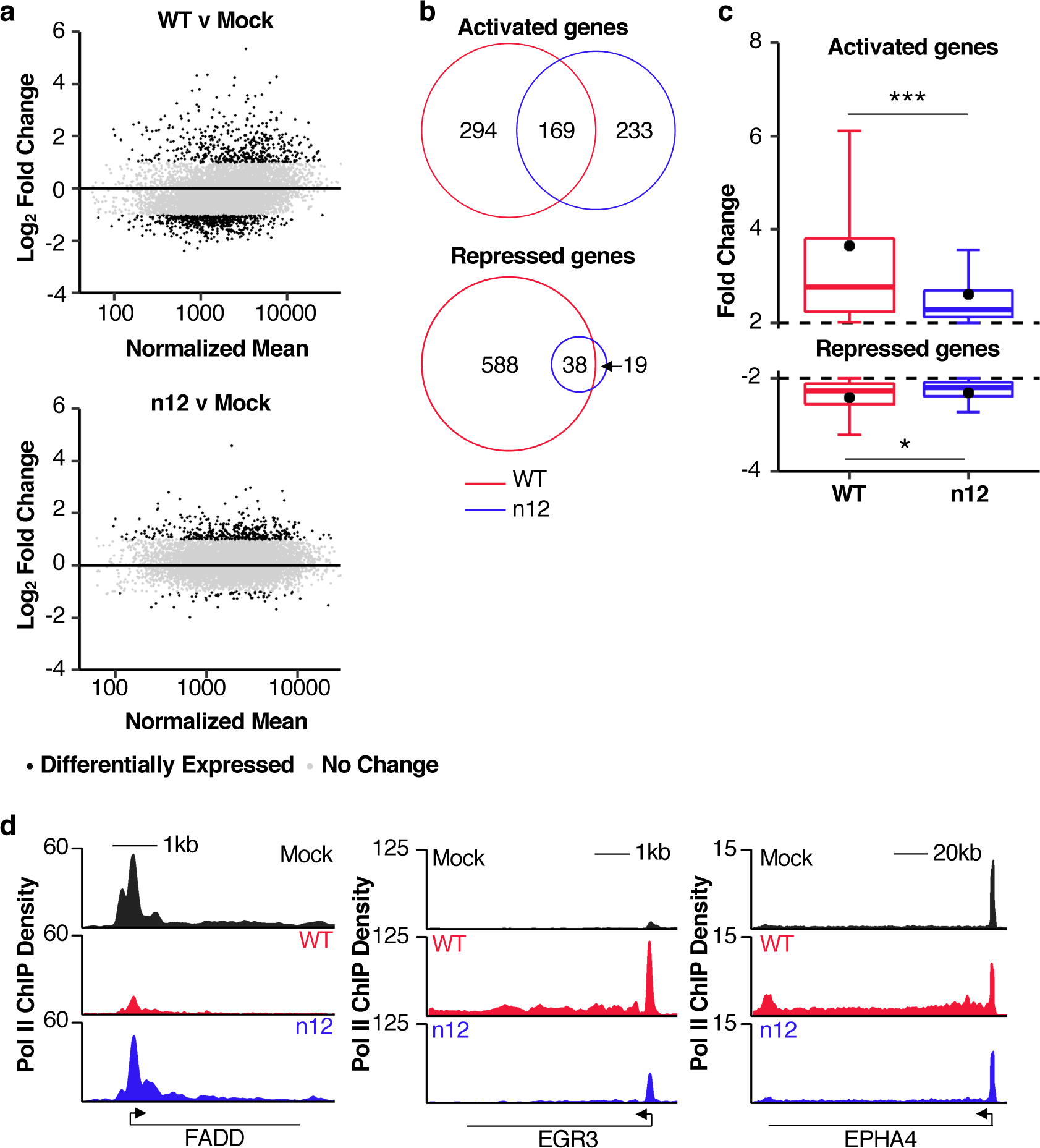
Removal of ICP4 dampens the increase in Pol II levels at activated host genes, and largely eliminates the decrease in Pol II levels at repressed genes. **(a)** Similar numbers of host genes show increases in Pol II occupancy after infection with the WT and n12 viruses, whereas very few host genes lose Pol II after n12 infection compared to WT infection. Shown are MA plots comparing Pol II levels in the bodies of cellular genes after WT infection (top) and n12 infection (bottom). For each gene, the log2 fold change in gene body RPK is plotted against the normalized mean across samples, as determined using DESeq2, to identify genes with significant changes in their Pol II gene body tags (+250 to the 3’ end). Black dots are genes with log2 fold change ≥1 or ≤-1 and Benjamini-adjusted p-values < 0.05. Gray dots represent genes that do not fit these criteria. 463 and 402 genes are activated upon WT and n12 infection, respectively. 626 and 57 genes are repressed upon WT and n12 infection, respectively. 7108 and 7886 genes are unchanged upon WT and n12 infection, respectively. **(b)** Overlap in the identities of the activated and repressed genes from both conditions are displayed using Euler diagrams. **(c)** The changes in Pol II occupancy on host cell gene bodies upon HSV-1 infection decrease in magnitude in the absence of ICP4. The quartiles of fold changes in gene body Pol II levels for activated genes (top) and repressed genes (bottom) are plotted for WT infection and n12 infection with the mean fold change represented as a black dot. P-values were determined by an unpaired Mann-Whitney test with *** representing a p-value < 2.2x10^-16^ and * representing a p-value < 0.05. Not shown are 20 and 19 outliers for WT and n12 activated genes, respectively, and 24 and 5 outliers for WT and n12 repressed genes, respectively. **(d)** Representative Pol II occupancy over three hosts genes: FADD (repressed by WT infection), EGR3 (activated by WT and n12 infection), and EPHA4 (activated by WT infection).

We next considered how the magnitude of changes in Pol II occupancy varied between WT and n12 infection. In Figure 2c, the fold changes in Pol II occupancy after infection for activated (top) and repressed genes (bottom) were plotted as box plots, with the mean fold change shown as a black dot. For both activated and repressed genes, the mean and median fold changes were smaller in n12 infection. Therefore, ICP4 is required for strong transcriptional reprogramming on the host genome regardless of whether transcription is activated or repressed. Examples of Pol II ChIP-seq traces for three representative genes are shown in Figure 2d. The FADD gene showed repressed Pol II levels during infection with the WT virus but not the n12 virus, EGR3 showed activated Pol II levels after infection with either WT or n12 virus, but the level of activation was reduced with the n12 virus, and EPHA4 was uniquely activated during WT infection. Together, the data in Figure 2 show that ICP4 is an important regulator of Pol II occupancy at both activated and repressed host cell mRNA genes during HSV-1 infection.

We performed gene ontology (GO) analysis to determine whether the genes with activated or repressed Pol II levels due to infection with either virus were enriched for specific biological processes. WT and n12 activated genes had a statistically significant enrichment for several GO-terms (Supplemental Figure 2), three of which involve Pol II transcriptional regulation and are shared between the two viruses. Repressed genes in WT infection are enriched in multiple metabolic pathways; the number of genes repressed by n12 infection was too small for meaningful GO analysis. We also evaluated enrichment in Hallmark gene sets, which are analogous to GO terms but represent specific biological states and pathways (Supplemental Figure 3). Several gene sets are enriched in both WT and n12 activated genes (e.g. TNFA signaling, p53 pathway, UV response), while gene sets enriched in the WT repressed genes included biological pathways such as cell cycle regulation and metabolism.

We also asked if core promoter elements were enriched in genes activated or repressed after WT or n12 infection using the CentriMo tool in MEME Suite. Of 9 core promoter elements (TATA box, GC-box, MED-1, XCPE1, BREu, BREd, INR, CCAAT-box, DPE), only the TATA box was significantly enriched in WT activated gene promoters (Supplementary Figure 4). No element was enriched in the promoters of WT repressed genes or any of the differentially expressed genes by n12 infection. We used the FIMO tool in MEME Suite to look for the consensus ICP4 binding motif in the region from -1000 through +100 relative to the TSS. There were only four instances of the ICP4 motif with an FDR < 0.05 in WT activated gene promoters, and only 5 instances in WT repressed gene promoters. These results suggest that ICP4-dependent effects on Pol II occupancy during HSV-1 infection do not require promoters to have ICP4 DNA motifs. We also found that no specific transcriptional activator or repressor binding motifs in the region from -1000 through +100 were enriched for genes with activated or repressed levels of Pol II after infection with WT virus.

### HSV-1 infection causes an increase in promotor proximal Pol II at activated genes independent of ICP4, and a decrease at repressed genes that depends on ICP4

Pol II ChIP-seq data show large peaks of Pol II occupancy just downstream from the TSS due to promoter proximal pausing. We evaluated how ICP4 impacts levels of promoter proximal paused Pol II occupancy on genes activated or repressed during WT HSV-1 infection using the ChIP-seq data from WT, n12, and mock infected cells. WT repressed genes experienced a large loss in promoter proximal Pol II during infection that was partially recovered in the absence of ICP4 (Figure 3a), consistent with prior literature [18]. Thus, the loss in promoter proximal Pol II due to HSV-1 infection is largely dependent upon ICP4. For genes activated by the WT virus, promoter proximal Pol II occupancy moderately increased after infection with either the WT or n12 virus when compared to mock (Figure 3b); therefore, the increase in promoter proximal Pol II at HSV-1 activated genes does not require ICP4.

**Figure 3.**
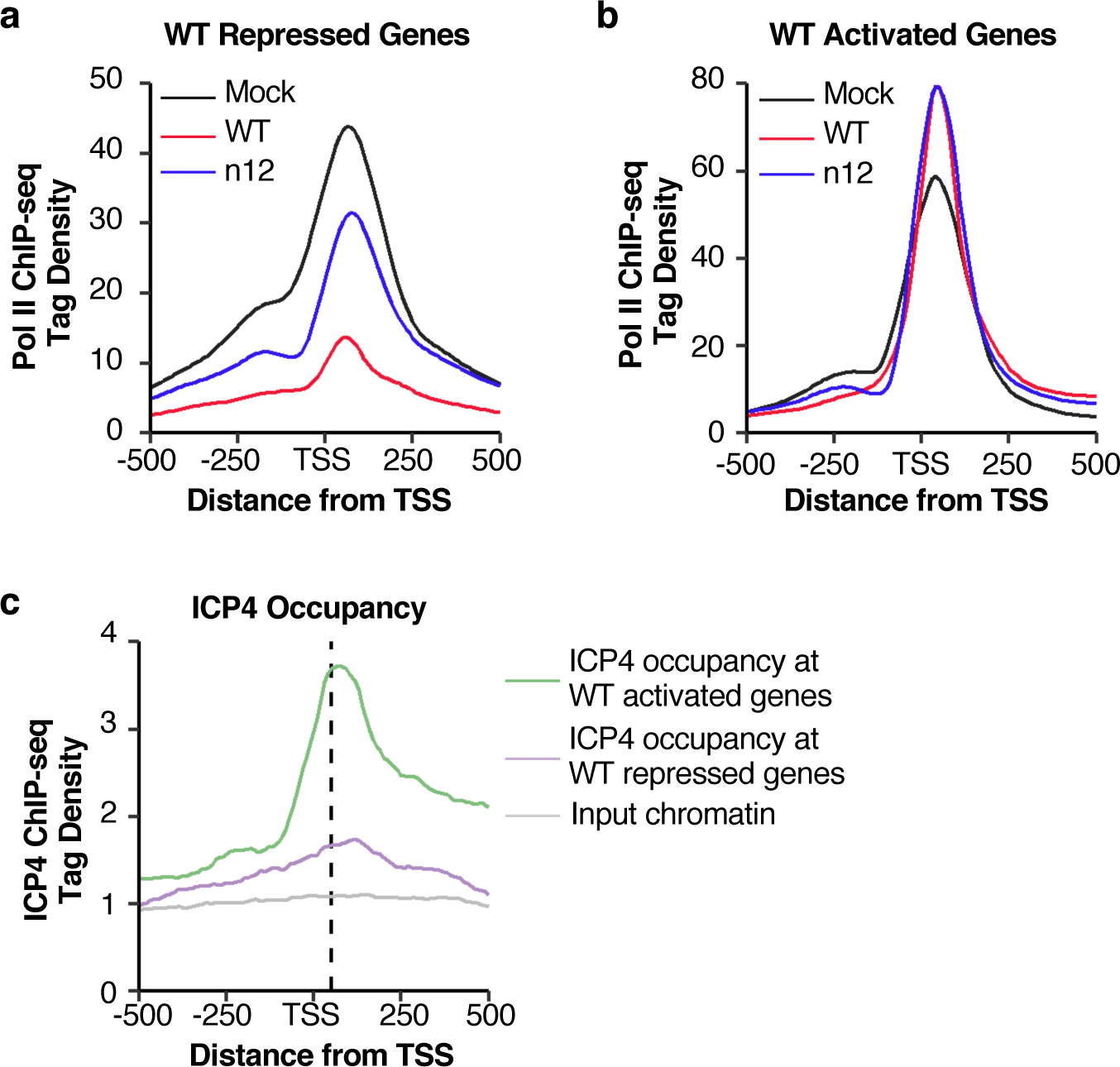
ICP4 modulates promoter proximal Pol II occupancy at cellular repressed genes, but not activated genes. **(a)** There is a substantial decrease in paused Pol II near the promoters of genes repressed by WT virus infection that depends on ICP4. For the 626 genes repressed after WT virus infection, Pol II ChIP-seq reads normalized to a constant read depth were binned in 10-bp bins and plotted from -500 to +500 with respect to the TSS. Reads from the mock, WT, and n12 conditions are shown. **(b)** Host genes activated after WT infection show a moderate increase in paused Pol II that is independent of ICP4. For the 463 genes activated after WT virus infection, Pol II ChIP-seq reads normalized to a constant read depth were binned in 10-bp bins and plotted from -500 to +500 with respect to the TSS. Reads from the mock, WT, and n12 conditions are shown. **(c)** ICP4 occupancy is higher near the promoters of genes activated by WT infection compared to those repressed by infection, and it correlates with Pol II occupancy. ICP4 ChIP-seq reads from a 4h infection with WT virus in MRC5 cells[18], normalized to a constant read depth, were binned in 10-bp bins and plotted from -500 to +500 with respect to the TSS. Reads mapping to the 463 genes activated (green) or 626 genes repressed by WT infection (purple), as determined by the Pol II ChIP-seq experiments, were plotted along with reads from input chromatin (gray). The input chromatin reads are the average of reads mapping to both the activated and repressed genes. The average position of the Pol II promoter proximal peak for the activated and repressed genes is shown as a black dashed line.

Near the start sites of genes, Pol II also transcribes in the opposite direction in a process known as divergent transcription [35], which is reflected by the shoulder of Pol II occupancy around -250 in our ChIP-seq data. Interestingly, at genes activated during HSV-1 infection, the divergent transcription peak was lost upon infection with WT virus and partially restored upon infection with the n12 virus (see Figure 3b), suggesting a role of ICP4 in regulating divergent transcription. To assess this, we calculated a divergent transcription ratio by dividing the reads in the upstream region (-300 to the TSS) by the reads in the downstream region (TSS to +300), to evaluate infection-induced changes in the ratio of divergent transcription to promoter proximal paused Pol II [36]. The data were log2 transformed and plotted as a cumulative distribution (Supplemental Figure 5). The divergent transcription ratios for WT and n12 infections are statistically different from the mock condition but are indistinguishable from each other. We conclude that HSV-1 infection alters divergent transcription in an ICP4-independent manner.

Prior ICP4 ChIP-seq experiments showed that ICP4 occupies the host genome at early time points of infection (2h and 4h) primarily around host gene promoters [18]. Although these data were collected in MRC5 cells, we wondered what ICP4 occupancy would look like at the genes we identified as activated or repressed due to WT infection. We made histograms of ICP4 occupancy around the TSSs of these genes using the publicly available ICP4 ChIP-seq datasets obtained after 4h of infection with WT HSV-1 (Figure 3c). ICP4 occupancy was highest on the activated genes (green line). At the repressed genes (purple line), ICP4 occupancy was less than 2-fold above the corresponding input chromatin (gray line). Notably, the peak of ICP4 occupancy aligns nicely with the shape and location of the promoter proximal peak in the Pol II ChIP-seq histograms (dashed black line). These data suggest that ICP4 is associated with promoter proximally paused Pol II on the host genome, particularly at activated genes, rather than binding directly to DNA.

### During HSV-1 infection, ICP4 facilitates the release of promoter proximally paused Pol II at activated host genes, but not repressed host genes

To further investigate the impact of HSV-1 infection and ICP4 on promoter proximal pausing, we calculated pausing indices for the set of genes activated during infection with the WT virus and plotted them as a cumulative distribution (Figure 4a). The pausing index at a gene is the ratio of the promoter proximal Pol II density (RPK TSS to +250) to the Pol II gene body density (RPK +250 to 3’-end). WT activated genes showed a strong shift to the left (i.e. a decrease) in the pausing index upon infection (red line compared to black line), suggesting the promoter proximal pause is a regulatory point during transcriptional activation due to HSV-1 infection. The curve shifts back towards the mock condition in n12 infected cells (blue line), indicating that ICP4 is important for controlling promoter proximal pausing at WT activated genes. We also evaluated the pausing index at the 294 genes that were dependent on ICP4 for activation during HSV-1 infection (see Figure 2b). The pausing indices at these genes strongly decreased upon infection with the WT virus but not during infection with the n12 virus (Supplementary Figure 6), which further supports a role for ICP4 in regulating promoter proximal pausing.

**Figure 4.**
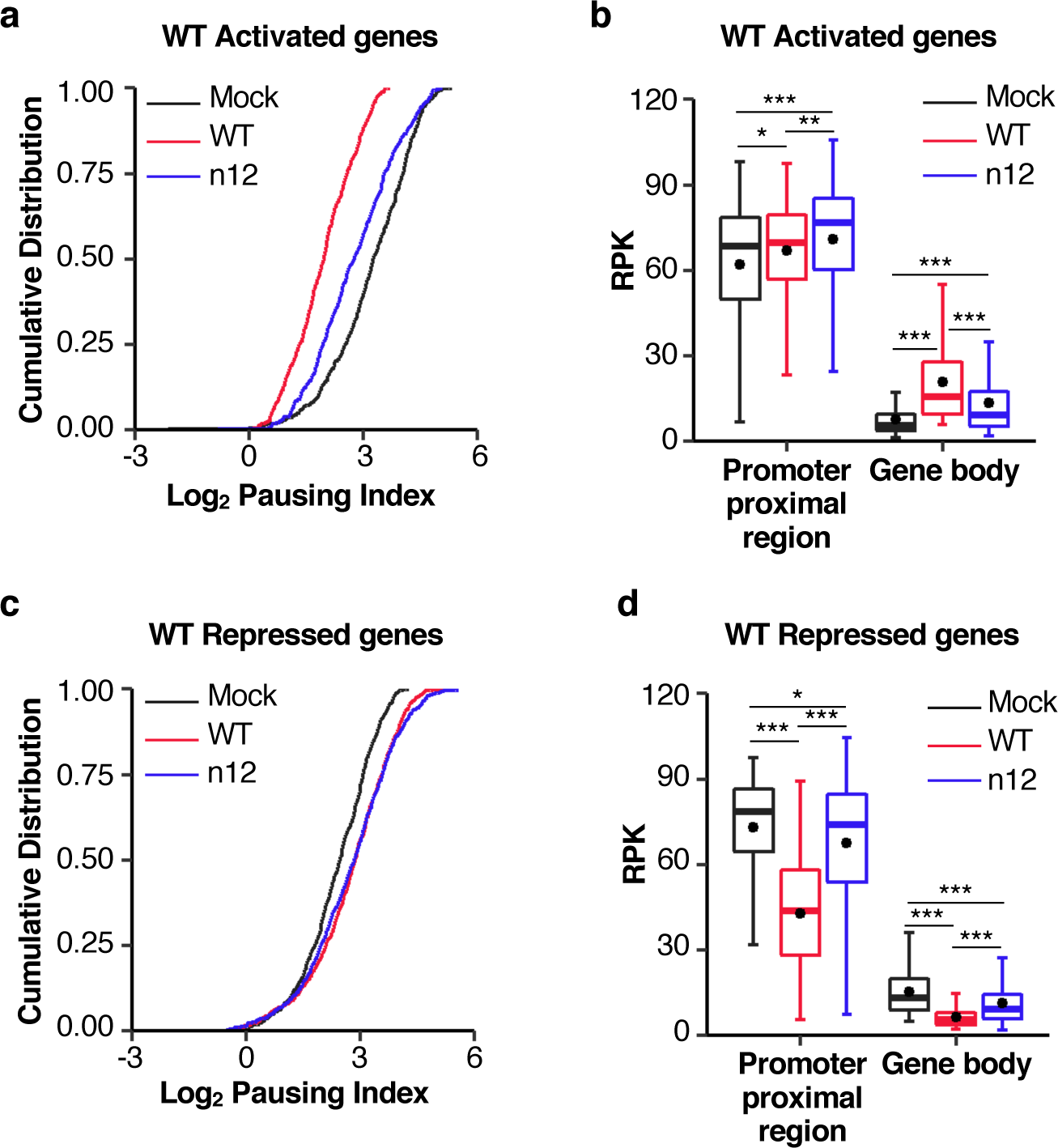
ICP4 regulates promoter proximal Pol II pausing at host cell genes activated by HSV-1 infection, but not at genes repressed by infection. **(a)** At cellular genes activated by the WT virus there is a strong decrease in the pausing index that is mostly recovered in the absence of ICP4. Plotted is a cumulative distribution of the log2 pausing indices (gene body RPK/promoter proximal RPK) for the 463 genes activated by WT virus infection, for the mock, WT, and n12 conditions. P-values were determined by using a two-sided Kolmogorov-Smirnov test: mock versus WT, p < 2.2x10^-16^; mock versus n12, p < 6.9x10^-9^; WT versus n12, p < 2.2x10^-16^. **(b)** The decrease in the pause index at host cell activated genes is primarily due to a large increase in Pol II occupancy in the gene bodies after infection. Shown are box plots comparing promoter proximal (+1 to +250) RPK and gene body (+250 to 3’-end) RPK at the 463 WT activated genes, across the three experimental conditions. P-values were determined by an unpaired Mann-Whitney test; * represents p-values < 0.050; ** represents p-values < 1.0x10^-10^; *** represents p-values < 2.2x10^-16^. **(c)** At cellular genes repressed by the WT virus there is a moderate increase in the pausing index that occurs independent of the presence of ICP4. Plotted is a cumulative distribution of the log2 pausing indices (gene body RPK/promoter proximal RPK) for the 626 genes repressed by WT virus infection, for the mock, WT, and n12 conditions. P-values were determined by using a two-sided Kolmogorov-Smirnov test: mock versus WT, p < 9.3x10^-10^; mock versus n12, p < 2.8x10^-9^; WT versus n12, p = 0.28. **(d)** The increase in the pause index at cellular repressed genes is dominated by a decrease in the gene body levels of Pol II after infection. Shown are box plots comparing promoter proximal (+1 to +250) RPK and gene body (+250 to 3’-end) RPK across the three experimental conditions for the set of WT repressed genes. P-values were determined by an unpaired Mann-Whitney test; * represents p-values < 0.05; *** represents p-values < 2.2 x10^-16^.

A decrease in pausing index can result from either a decrease in Pol II occupancy within the promoter proximal region (+1 through +250), an increase in Pol II gene body occupancy, or a combination of the two. For WT activated genes, we had observed a moderate increase in the promoter proximal peak upon infection (see Figure 3a), suggesting that Pol II gene body occupancy must have substantially increased during infection to result in the large decrease in the pausing index we observed at these genes. To investigate this, we generated box plots of the Pol II density in the promoter region (RPK TSS to +250) and across gene bodies (RPK +250 to 3’-end) for the WT activated genes (Figure 4b). As expected, the Pol II occupancy in the gene bodies substantially increased during WT infection compared to the mock condition, and this increase was significantly reduced in n12 infection. The differences between the conditions in the promoter region are statistically significant, however, the magnitude of the differences is much smaller. We calculated the ratio of the mean RPK in the promoter region to the mean RPK in the gene body for all three conditions. In mock infected cells, this ratio is 8.11; it decreases to 3.22 in WT infected cells and is 5.29 for n12 infected cells, agreeing with the trends in the pausing index observed in Figure 4a. Together these data show that ICP4 promotes the transition of promotor proximally paused Pol II into transcriptional elongation at genes activated upon HSV-1 infection.

We performed a similar analysis of promoter proximal pausing at genes repressed after infection with the WT virus. We calculated the pausing indices for these genes across all three conditions (Figure 4c). In contrast to the activated genes, we found that at repressed genes infection caused a statistically significant increase in the pausing index relative to mock infected cells (red line compared to black), which did not require ICP4 (blue line). The magnitude of the increase in pause index was much smaller at the repressed genes compared to the magnitude of the decrease observed for activated genes. A box plot was created for the Pol II density in the promoter and gene body regions for the WT repressed genes (Figure 4d). As expected, the Pol II occupancy decreased in the promoter region due to WT infection (see Figure 3b); however, a larger decrease was observed in the gene body region (2.4-fold, using the mean RPK) than in the promoter proximal region (1.7-fold, using the mean RPK), comparing mock to WT data. This explains the increase in the pause index we observed at WT repressed genes. Decreases in Pol II occupancy in both the promoter and gene body regions were attenuated in the absence of ICP4, with the ratio between the two regions similar for the WT and n12 virus. Therefore, ICP4 contributes to the overall loss of Pol II on both regions of host genes repressed during HSV-1 infection but does not change the pause index.

Given our data suggesting that ICP4 promotes pause release at activated genes, we asked whether infection with the WT and n12 virus altered the occupancy of the pausing factors NELF and DSIF at the promoter of a representative activated gene. We performed ChIP assays with antibodies targeting the NELF-A and Spt5 subunits of NELF and DSIF, respectively, as well as with the total Pol II antibody in mock and WT or n12 infected cells. The promoter proximal region of EPHA4 was amplified from the ChIP eluates and compared to a standard curve of the input chromatin to determine the percent IP. Since both NELF and DSIF bind to Pol II, and Pol II levels change upon infection, we normalized the NELF-A and Spt5 occupancy by the Pol II occupancy. As shown in Figure 5, both NELF-A and Spt5 occupancy decreased relative to Pol II occupancy in the EPHA4 promoter proximal region upon WT infection, and increased back to mock levels in n12 infected cells. These data show that ICP4 causes a decrease in pausing factor occupancy at a gene activated by HSV-1 infection. Together our data support a model where ICP4 promotes Pol II pause release by controlling NELF and DSIF occupancy.

**Figure 5.**
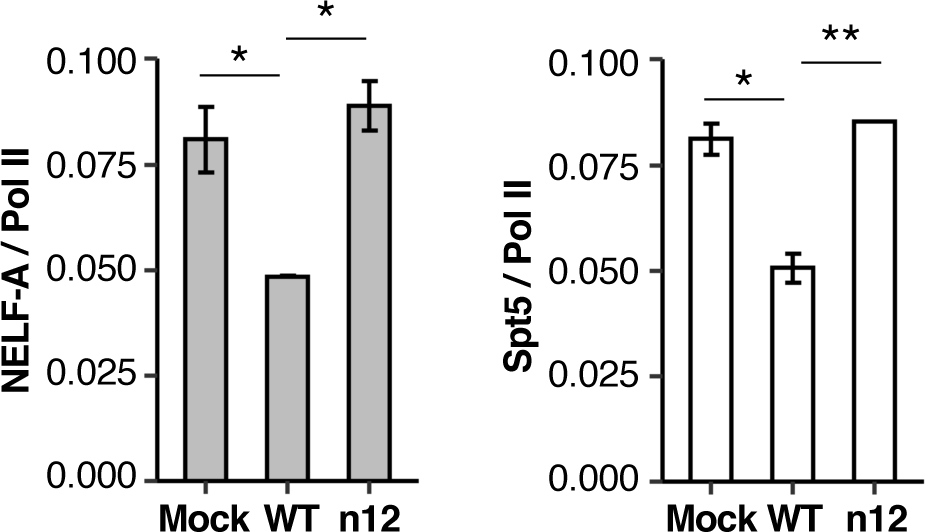
HSV-1 infection decreases the occupancy of NELF-A and Spt5 with respect to Pol II at the promoter of an activated gene in an ICP4 dependent manner. Plotted is the average NELF-A occupancy (gray bars) or Spt5 occupancy (white bars) as a ratio of Pol II occupancy in the region +1 to +150 relative to the TSS for the WT activated gene EPHA4. The ratios were calculated using the percent IPs obtained with each antibody in ChIP assays. Error bars are the range of two biological replicates. The error bars for WT infection and n12 infection for NELF-A and Spt5, respectively, are too small to visualize. P-values were determined by an unpaired student *t* test; * represents p-values < 0.05; ** represents p-values < 0.01.

### Exogenous expression of ICP4 activates genes but is sensitive to the chromatin environment

We were interested to see if ICP4 could modulate Pol II transcription in the absence of infection. We transfected HEK 293 cells with either a plasmid to express mCherry-tagged ICP4 (mChy-ICP4) or with a plasmid to express mCherry with 3 nuclear localization sequences (mChy-NLS) as a control, then 24h later sorted and collected mChy-positive cells. Total RNA was extracted, and RT-qPCR was performed to detect specific mRNA transcripts from genes that showed repressed or activated Pol II levels in response to HSV-1 infection in our ChIP-seq data. For two genes repressed in an ICP4-dependent manner during WT HSV-1 infection, FADD and INCENP, we observed no significant change in their mRNA levels due to overexpression of mChy-ICP4 (Figure 6a, first two bars); hence, ICP4 is not sufficient to repress these genes in the absence of infection. For four genes activated during WT HSV-1 infection (IGF2, EPHA4, VEGFA, and EGR3), levels of VEGFA and EGR3 increased upon overexpression of mChy-ICP4, whereas levels of the other two mRNAs did not change (Figure 6a). We again used the FIMO tool to look for the presence of ICP4 motifs in the promoters of these 7 genes and found none. These data suggest that ICP4 is sufficient to activate some genes in the absence of infection, independent of an ICP4 binding motif, and that repression of two genes by ICP4 requires infection.

**Figure 6.**
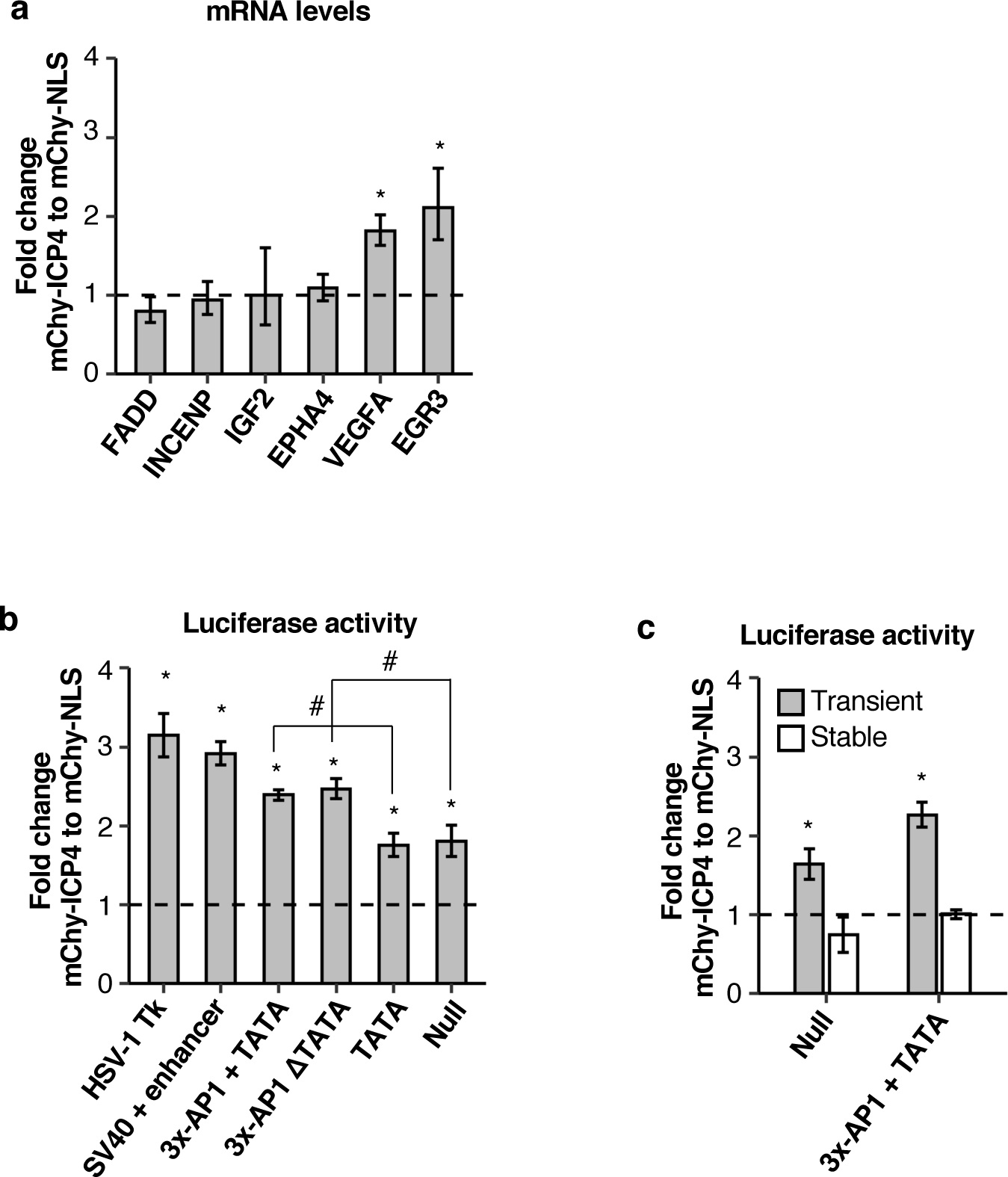
ICP4 is sufficient to activate gene expression in the absence of infection under some conditions. **(a)** ICP4 selectively activates cellular genes in the absence of infection. Cells expressing mChy-ICP4 or mChy-NLS were collected via FACS, and fold changes in endogenous mRNA levels due to ICP4 were calculated using qRT-PCR and the ΔΔC_T_ method after first normalizing transcript levels to 18S rRNA levels. Error bars are the standard deviation of three biological replicates. P-values were determined by a one sample *t* test comparing the ΔΔC_T_ value to a mean value of zero; * represents p-values < 0.05. **(b)** ICP4 activates luciferase expression from transiently transfected reporter plasmids independent of their promoter elements. Plotted is the fold change in luciferase expression due to overexpression of mChy-ICP4 compared to the negative control mChy-NLS for the reporter plasmids indicated. Error bars are the standard deviation of three biological replicates. P-values were determined by a one sample *t* test comparing mean fold changes to 1.0; for all plasmids tested p < 0.05, denoted by *. P-values were also calculated using an unpaired t-test to evaluate differences between the constructs containing 3xAP-1 and/or TATA elements. Only the differences between 3xAP-1 versus TATA and 3xAP-1ΔTATA versus null were significant (p < 0.005), indicated by #. **(c)** ICP4 cannot activate luciferase from episomally associated plasmids. Plotted is the fold change in luciferase expression after overexpression of mChy-ICP4 compared to the mChy-NLS control in cells either transiently expressing luciferase with the indicated promoter (gray bars) or stably expressing luciferase with the indicated promoter (white bars). Error bars are the standard deviation of three biological replicates. P-values were determined by a one sample *t* test comparing the mean fold changes to 1.0; * indicates p < 0.05.

We asked if activation by exogenous ICP4 was dependent on promoter composition by co-transfecting mChy-ICP4 with a series of luciferase reporter plasmids. We first tested a reporter plasmid that contains the HSV-1 Tk promoter, which is from a viral gene activated early during HSV-1 infection in an ICP4 dependent manner, and for comparison a reporter plasmid that contains the SV40 promoter/enhancer. As shown in Figure 6b, left two bars, expression of mChy-ICP4 caused a ∼3.2-fold increase in luciferase expression from the HSV-1 Tk promoter relative to mChy-NLS. A similar level of activation was observed with the SV40 promoter, even though this promoter does not have ICP4 binding sites. We next tested a promoter that contains three binding sites for the host AP-1 activator proteins and a TATA box upstream of the firefly luciferase gene. mChy-ICP4 activated luciferase expression from this plasmid 2.4-fold compared to mChy (Figure 6b third bar); this activation did not require the TATA box (fourth bar, TATA box mutated from TATATAAT to TAGCTAGC). Lastly, we tested whether ICP4 would activate luciferase expression from plasmids that contain just the TATA box or with no known promoter elements (null). With these reporters mChy-ICP4 activated luciferase expression (Figure 6b, right two bars), however, to a lower degree than the analogous reporter plasmids that contained the three AP-1 sites. Hence, activation by ICP4 is not dependent on the presence of the TATA-box or AP-1 DNA elements.

We next investigated whether activation by exogenous ICP4 was sensitive to the chromatin state of the gene. To that end, we created a stable cell line harboring an episomally associated luciferase expression plasmid. Episomal plasmids are chromatinized and more closely resemble euchromatin on the genome than transiently transfected plasmids [37, 38]. We used two different plasmids: one with no known promoter elements (null) and another with three AP-1 sites plus a TATA box. mChy-ICP4 was unable to activate luciferase expression from plasmids in the stable cell lines (Figure 6c, white bars), while mChy-ICP4 activated luciferase expression during transient transfection of each plasmid (Figure 6c, gray bars). These data indicate that the chromatin state of a gene may affect whether a gene is activated by ICP4.

## Discussion

Here we investigated the role of the HSV-1 ICP4 protein in controlling transcription of host genes during viral infection. Pol II ChIP-seq before and after infection with WT and n12 virus enabled us to quantify changes in the amounts and locations of Pol II bound on the human genome during infection. We found that ICP4 was required for the repression of gene body Pol II levels at the majority of host genes repressed during infection with the WT virus. ICP4 was also required for ∼60% of the WT infection-induced activation of Pol II levels on host genes, with ∼200 genes uniquely activated during infection in the absence of ICP4 (n12 virus). We found that ICP4 promoted the release of paused Pol II on activated genes and was required for reduced NELF-A and Spt5 occupancy on a representative activated gene, suggesting a mechanism by which ICP4 reduces pausing to promote elongation complex formation. By contrast, repressed genes showed a slight increase in pausing independent of ICP4, suggesting that ICP4 inhibits recruitment of Pol II at repressed genes during infection. Exogenous expression of ICP4 in the absence of infection showed it is sufficient to activate transcription in some cases but cannot itself repress transcription. ICP4 could activate luciferase expression from transiently transfected plasmids irrespective of promoter composition; however, ICP4 could not activate luciferase expression on stable, episomally associated plasmids. We propose that ICP4 has dual-activities capable of both activation and repression, the former of which is sensitive to the chromatin state of the gene, while the latter relies on the presence of other viral proteins or cellular changes due to viral infection.

We found that within 4h of HSV-1 infection of human cells there are dramatic changes to the host cell mRNA transcriptional program, with hundreds of genes losing Pol II occupancy and hundreds of others gaining Pol II. This is largely consistent with the transcriptional reprogramming observed in prior studies that also studied the effect of HSV-1 infection on host cell transcription in human cells looking at nascent mRNA production via PRO-seq or 4SU-seq rather than using Pol II ChIP-seq [10,11,13,39,40]. The effect of HSV-1 infection on Pol II occupancy in human cells is significantly different from that seen in mouse cells where only the reduction of Pol II occupancy was observed after 4h of infection with little to no activation [12], suggesting that the activation of Pol II transcription is a signature specific to the host species. Prior studies have also documented that HSV-1 infection disrupts transcription termination by Pol II as well as splicing [13,17,40,41]. Strong termination and splicing defects occur on average later during infection than the 4h time point investigated here. Nonetheless, the termination defect was considered in our bioinformatic analyses of Pol II ChIP-seq reads (see Methods section) to ensure genes were not falsely designated as activated due to failed termination of a neighboring gene.

We observed that ICP4 plays a fundamental role in decreasing Pol II occupancy on nearly all genes that were repressed during HSV-1 infection. This is consistent with a recent study implicating ICP4 as the viral protein responsible for reducing Ser5-P modified Pol II occupancy at the promoter proximal region of repressed genes [18]. Here we used an antibody that targets total Pol II irrespective of the phosphorylation state of the Pol II CTD, thereby allowing us to investigate viral-induced Pol II occupancy changes in the entire gene body, which closely reflects ongoing transcription [23]. The use of a total Pol II antibody revealed that the promoter proximal peak at genes repressed by HSV-1 infection decreased, which was largely alleviated during infection with the n12 virus (Figure 3a). Further analyses revealed that at repressed genes the loss in gene body Pol II was greater than the loss observed at promoter proximal peaks, leading to an increased pause index (Figures 4c and 4d). Together our data support a model that at host genes repressed during infection, ICP4 acts by blocking either recruitment of Pol II to promoters and/or transcription initiation.

Although ICP4 was required for the loss in Pol II at nearly all of the genes considered repressed, the levels of Pol II at the promoters of these genes did not fully return to the mock-infected levels when ICP4 was eliminated and the increase in the pausing index did not require ICP4. This indicates that although ICP4 is the primary viral protein functioning to repress host cell transcription, additional proteins (viral or cellular) are likely involved. Many of the other IE proteins have roles in transcriptional regulation, including interactions with host machinery. For example, ICP27 co-immunoprecipitates with Pol II [16] and has recently been identified to control transcription termination of host genes [13, 17]; ICP0 blocks the association of HDAC1 with REST/CoREST complexes that are known to influence transcription [2, 42]; ICP22 interacts with PTEF-b and FACT, which are also known transcriptional regulators [43, 44], as well as influences the phosphorylation state of the CTD [45]. Furthermore, the presence of viral DNA could act as a sink for transcription machinery contributing to repression of host genome transcription. Replication compartment formation upon the expression of Early viral proteins assists in sequestering transcription machinery near viral genomes [46]. After viral replication, there is a 100-fold higher density of ICP4 motifs present on the viral genome than the host genome, which has been proposed to sequester transcription machinery through mass action [18]. Either or both of these models could contribute to a general decrease in Pol II occupancy on the host cell genome.

Our experiments showed that ICP4 plays a role in activating host gene transcription during HSV-1 infection. This is consistent with its known role in activating transcription of genes on the HSV-1 genome [4]; however, our observations indicate that ICP4’s effects on host transcription are somewhat gene specific. ICP4 plays a clear role in activating ∼300 host genes during infection, but ∼170 are still activated in the absence of ICP4. Of the genes whose activation depends upon ICP4, we did not identify an enrichment in ICP4 binding motifs in the promoter regions of these genes, suggesting a mechanism that does not involve ICP4 directly binding the genome. Surprisingly, infection with the n12 virus caused 233 genes to become activated that were not activated by infection with the WT virus. It is possible that the transcriptional activation of these genes during WT infection is normally dampened by ICP4; however, during infection with the n12 virus, these genes are activated. Given that GO analysis revealed that many of the WT and n12 activated genes are involved in transcriptional regulation, it is possible that activation of many genes is a downstream effect from upregulating transcriptional regulatory proteins.

To understand the mechanism of transcriptional activation by ICP4 at host genes during infection, we analyzed promoter proximal pausing. During infection, the pausing index decreased substantially at WT activated genes, which was dependent on ICP4 (Figure 4a). Further investigation showed that ICP4 decreased the pause index primarily by increasing gene body Pol II occupancy. Together these data are consistent with a model in which promoter proximal pausing occurs at activated genes, but ICP4 decreases the residence time of the paused complexes. This would increase the release of paused Pol II into elongation complexes, resulting in overall gene activation. It is possible that ICP4 promotes the release of paused Pol II by decreasing the residence of pausing factors. Consistent with this, ChIP-qPCR assays against the pausing factors NELF-A and Spt5 at a candidate gene revealed their occupancy was decreased in the promoter proximal region during WT infection, which was dependent upon the presence of ICP4 (Figure 5). The HSV-1 protein ICP22 interacts directly with the CyclinT1 subunit of PTEF-b to inhibit phosphorylation of Ser2-P on the Pol II CTD [14, 43]; however this interaction does not affect the recruitment of PTEF-b to host genes [43]. To date, studies have not revealed the global consequences of HSV-1 infection on NELF and DSIF occupancy at paused Pol II on the host genome. Our ChIP experiments suggest that ICP4 could play a critical role in globally regulating their association with Pol II at activated host genes during infection.

To understand whether ICP4 could control Pol II transcription in the absence of infection, we performed exogenous expression assays with mChy-ICP4. We discovered that the effects of exogenously expressed ICP4 did not mirror the ICP4-dependent effects we saw with WT infection. Of the 6 endogenous genes we studied, the mRNA levels from only 2 increased in response to ICP4 overexpression and none were repressed. The observation that ICP4 is incapable of repressing transcription in the absence of infection surprised us since we and others [18] found that loss of Pol II on the genome after infection is strongly dependent upon ICP4. This supports the model that infection-dependent factors in addition to ICP4 are required to repress host transcription during infection. Only VEGFA and EGR3 were activated by exogenous ICP4. Interestingly, others have reported that VEGFA transcription is dependent on ICP4 for activation during HSV-1 infection [47]. The authors hypothesize that the VEGFA promoter is GC-rich and resembles viral promoters allowing VEGFA to escape repression upon infection.

In our transient transfection luciferase reporter assays we observed universal activation of luciferase expression by ICP4 regardless of the identity of the promoter or the presence of a TATA box, although the level of activation varied between constructs. By contrast, during HSV-1 infection, the TATA-box was enriched at activated genes, the majority of which were ICP4-dependent. Our reporter assays did suggest that ICP4-dependent activation in the absence of infection is sensitive to the chromatin state of the gene since activation was only observed from transiently transfected plasmids as opposed to stably maintained episomal DNA know to be better chromatinized [37, 38]. Exogenous expression of ICP4 in the absence of infection increases the free pool of histone H3.1 [48]; however, how this interplay between ICP4 and chromatin relates to transcriptional activity is unclear. During infection, ICP4 binding the host genome correlated with euchromatic histone marks and accessible chromatin without changing the euchromatic markers that were present prior to infection [18]. The precise relationship between chromatin and the ability of ICP4 to drive activation or repression of transcription activity requires further investigation.

Together our data provide evidence that ICP4 acts as a transcriptional activator and repressor on the host genome during HSV-1 infection. ICP4 represses transcription by blocking Pol II recruitment to promoters and/or initiation, while activates transcription by facilitating Pol II pause release into productive elongation. Activation by ICP4 is likely facilitated by the chromatin state of the gene, however the mechanism is still unclear. Moreover, transcriptional repression by ICP4 requires both infection and additional unidentified factors. Ongoing studies will continue to reveal the novel regulatory relationships between host and virus.

## Materials and Methods

### Cells and viruses

HEK 293 cells (American Type Culture Collection, ATCC) were cultured in Dulbecco’s modified Eagle’s medium (DMEM) containing 10% fetal bovine serum (FBS), 100 U/ml penicillin and 100 mg/ml streptomycin at 37°C in 5% CO_2_. HEK 293 cells were authenticated by STR profiling and regularly checked for mycoplasma infections. The WT virus (HSV-1 KOS strain, provided by D. Bloom) was prepared and titered in Vero (African green monkey kidney) cells (ATCC); n12 stocks (provided by N. DeLuca) were prepared and titered in a Vero-based ICP4-complementing cell line [33]. HEK 293 cells were infected at ∼80% confluency with either the WT virus at a multiplicity of infection of 10 pfu/cell or n12 virus to produce a viral load equivalent to the WT infection, as determined using qPCR of isolated viral genomes from infected cells. After 1h, the media was removed and replaced with fresh, prewarmed media for the remaining 3h prior to harvesting.

### Chromatin immunoprecipitation assays

After 4h of infection with WT virus, n12 virus, or media (mock-infected samples), cells (∼2.5 × 10^7^) were crosslinked with 1% formaldehyde for 10 min at room temperature then quenched with glycine (0.125M) for 5 min. Nuclei were isolated by resuspending cells in lysis buffer (80 μl/million cells; 4 mM MgCl2, 10 mM Tris-HCl (pH 7.4), 10 mM NaCl, 0.5% NP-40, 0.4 mM phenylmethylsulfonyl fluoride (PMSF), and protease inhibitors (Complete cocktail tablets; Roche), incubating on ice for 6 min, spinning at low speed, then washing the nuclei once in lysis buffer. The nuclei were resuspended in resuspension buffer (30 μl/million cells; 15 mM Tris pH 7.9, 10 mM EDTA, 0.4 mM PMSF, 1% SDS, and protease inhibitors) and nutated for 10 min at 4°C. Dilution buffer was then added (60 μl/million cells; 15 mM Tris (pH 7.9), 150 mM NaCl, 1 mM EDTA, 1% Triton X-100, 0.4 M PMSF, protease inhibitors) and samples were sonicated using a Diagenode Bioruptor on the medium setting for a total of 30 min (30 sec, 30 sec off). Samples were clarified by centrifugation at high speed and the supernatants were used for immunoprecipitation. Pol II ChIP-seq experiments were performed using ∼12 million cells per biological replicate, as follows. ChIP-qPCR experiments were performed in biological replicate using 100 μl of sheared chromatin per IP using the total Pol II antibody, NELF-A antibody (sc-365004 G-11), or Spt5 antibody (sc-133217x D-3). Solubilized chromatin was precleared using protein A/G beads (Santa Cruz Biotechnology) equilibrated in buffer D (15 mM Tris (pH 7.9), 150 mM NaCl, 1 mM EDTA, 0.5% NP-40, 0.4 mM PMSF) by nutation for 2h at 4°C. The D8L4Y antibody targeting the N-terminal region of Rpb1 (Cell Signaling Technology) was added to precleared chromatin and samples were nutated overnight at 4°C. Protein A/G beads were equilibrated in buffer D, pre-blocked with total yeast RNA (0.4 mg/ml) and bovine serum albumin (BSA, 0.5 mg/ml), and added to chromatin-antibody complexes, followed by nutation for 3h at 4°C. Beads were washed sequentially with low-salt buffer (20 mM Tris (pH 7.9), 150 mM NaCl, 2 mM EDTA, 1% Triton X-100, 0.1% SDS), high-salt buffer (20 mM Tris (pH 7.9), 500 mM NaCl, 2 mM EDTA, 1% Triton X-100, 0.1% SDS), LiCl buffer (20 mM Tris (pH 7.9), 1 mM EDTA, 1% deoxycholate, 1% NP-40, 250 mM LiCl), and twice with TE buffer (10 mM Tris (pH 7.9), 1 mM EDTA). Chromatin was eluted by nutation for 1h at 37°C with 1% SDS, 50 mM Tris Base (pH 7.9), and 10 mM EDTA. Supernatants were moved to a new tube and NaCl was added to a final concentration of 200 mM. Cross-links were reversed overnight by incubation at 65°C. Samples were diluted 1:1 with water and treated with Proteinase K (80 μg) for 1h at 55°C, phenol/chloroform/isoamyl alcohol extracted (pH 8.0), and ethanol precipitated. qPCR was performed using the Luna Universal qPCR Master Mix (NEB) in technical duplicate with a StepOnePlus Real-Time PCR system (Applied Biosystems) with SYBR Green detection. The percent IP was calculated by comparing the signal from the ChIP eluates to a standard curve of input chromatin. Primer sequences used are found in Supplemental Figure 7.

### High-throughput sequencing and read mapping to the human genome

Libraries for Illumina sequencing were prepared from the Pol II ChIP eluates by the University of Colorado Anschutz Genomics and Microarray Core facility using the Tecan Ovation Ultralow DNA Library System. Sequencing was performed using an Illumina NovaSEQ6000, obtaining 2 × 150 bp reads. Total reads were as follows: mock replicate 1, 156,037,414; mock replicate 2, 146,116,674; mock input, 139,115,086; WT replicate 1, 158,551,068; WT replicate 2, 220,866,112; WT input, 134,363,468; n12 replicate 1, 131,051,270; n12 replicate 2, 185,062,944; n12 input, 172,126,626. The sequencing quality was assessed using FastQC. Reads were mapped to the human genome using the hg19 assembly with Bowtie 1.2.2 [49] in the -v alignment mode allowing one mismatch per read (-v1), mapping the first 50 nucleotides (--trim3 101), and only reporting reads that aligned to a unique position (-m1). With these settings, 64-81% of reads mapped to the human genome for ChIP samples and 43-53% mapped to the human genome for input samples.

### Computational analysis of mapped reads

Meta-analysis and per-gene quantification of mapped reads were performed using the suites of tools available in HOMER v4.9.1 (Hypergeometric Optimization of Motif EnRichment) [50] and Bedtools (v2.27.1) [51]. Tag directories were created in HOMER from the .map files generated by Bowtie. Tag directories were converted to BED file format for use in Bedtools and bigwig file format for visualization in R. UCSC RefSeq genes (75,683) from the UCSC Table Browser were filtered to remove non-coding RNAs and genes less than 1 kb in length. Our analyses considered one isoform per gene, chosen by the highest promoter proximal peak occupancy (TSS to +250), then highest termination peak occupancy (3’-end to +500), and then highest whole gene occupancy (TSS to 3’-end) using the Bedtools Coverage command, leaving 19,025 unique mRNA genes. Using this list of genes, the whole gene tag counts per biological replicate were compared against one another using linear regression, Pearson coefficients, and principal component analysis (see Supplementary Figure 1). After it was determined that the replicates were similar, the reads were pooled together for most downstream analyses.

Pol II ChIP-seq reads across gene bodies (+250 to 3’-end) were summed using the Bedtools coverage command for each biological replicate and the input chromatin. Then DESeq2 (v1.26.0) [52] was used to identify genes in which the gene body Pol II occupancy levels changed significantly (fold change ≥ 2; p-adj value ≤ 0.05) by comparing the WT or n12 infected samples to the mock condition. Of the differentially expressed genes, only those with Pol II levels higher than input chromatin (average gene body RPK between replicates ≥ 2-fold over the average input gene body RPK for all genes within a condition) were considered. HSV-1 infection causes a disruption of transcription termination [13, 40]; although most severe at later time points of infection, we filtered our lists of genes transcribed after infection to remove those with Pol II occupancy due to failed transcription termination of a nearby gene. First, we required that each gene had a promoter proximal peak (PPP), defined as having at least 23 RPK from the TSS to +250. In particular for activated genes, we required the reads from the TSS to +2kb to be at least 1.1-fold higher than the reads from -3kb to -1kb relative to the TSS. In addition, we required that the PPP RPK be at least 1.75-fold higher than the RPK from -3kb to -1kb upstream of the TSS. Genes around these thresholds were viewed individually on the UCSC Genome Browser as a verification. Altogether, 184 activated genes and 111 repressed genes were removed from WT differentially expressed genes, and 84 activated genes and 34 repressed genes were removed from n12 differentially expressed genes. To generate histograms, tag counts surrounding the TSS were binned using the annotatePeaks.pl command in HOMER using BED files for lists of genes. Gene ontology analysis was performed using the Functional Annotation tool in DAVID Bioinformatics Resources 6.8 [53] with the gene ontology categories GOTERM_BP_DIRECT (activated genes) or GOTERM_BP_ALL (repressed genes). Enriched terms had a Benjamini corrected p-value of < 0.05. Hallmark gene set analysis was performed using the Molecular Signatures Database (MSigDB) [54, 55]. Heatmaps of Pol II tag counts were generated using Cluster 3.0 and plotted in TreeView v1.1.6 [56]. Pol II occupancy traces were generated with BigWig files using the Gviz [57] and rtracklayer [58] libraries in R v4.0.3. All other plots were generated using the ggplot2 library in R. Motif analysis was performed using the HOMER suite as well as CentriMo and FIMO tools within MEME suite [59]. The ICP4 motif file used in FIMO was created using the iupac2meme tool in MEMEsuite using the sequence DTSGKBDTBNHSG.

### Transfection assays, luciferase assays, and cell sorting

Plasmids transfected included the psiCHECK2 vector (Promega), 3xAP-1+TATA pGL3 plasmid, 3xAP-1ΔTATA pGL3 plasmid, TATA only pGL3 plasmid, pRL-null plasmid (Promega), null-PREP4 (obtained from Addgene, plasmid #124892) [60], and 3xAP-1_pREP4. The 3x-AP-1ΔTATA plasmid was generated using quick change mutagenesis on the 3x-AP-1 pGL3 plasmid to mutate TATATAAT to TAGCTAGC. The 3xAP-1_pREP4 plasmid was generated using a gblock (IDT) that contained the promoter sequence from the 3x-AP-1 plasmid with Kpn1 and HindIII sites added to drop into the null pREP4. The linker sequences between the AP-1 sites were mutated to decomplex the gblock sequence for higher yield during oligo synthesis. The mChy-ICP4 plasmid was a gift from the L. Schang lab [48]. The mChy-NLS plasmid was obtained from Addgene (plasmid #39319) [61].

For transfections, HEK 293 cells were plated at ∼500,000 cells per well of a 6-well plate. 48h after plating, cells were co-transfected with 250 ng of luciferase reporter plasmid and 1 µg of either mChy-ICP4 or mChy-NLS using Lipofectamine 3000 reagents (ThermoFisher) per the manufacturer’s recommendation. After 24h, cells were harvested and lysed in cell culture lysis buffer (Promega). Luminescence was measured using the Dual-Luciferase Reporter Assay System (Promega) for psiCHECK2 transfections, *Renilla* Luciferase Assay System (Promega) for null promoter *Renilla* transfections, and Luciferase Assay System (Promega) for all other transfections. For all transfections, luciferase signals in mChy-ICP4 transfections were normalized to luciferase signal in the mChy-NLS transfections. For pREP4 plasmids, 750 ng of pREP4 plasmid was transfected. For stable expression of pREP4 plasmids, cells were transfected with 750 ng plasmid and selected for with 250 µg/mL hygromycin B (Mirus Bio) for 14 days; selection was then maintained with 125 µg/mL drug. Transfections with the mChy-ICP4 and mChy-NLS plasmids into these cells were done in the absence of drug but after stable selection was achieved.

To observe the impact of exogenous ICP4 on endogenous genes in the absence of infection, HEK 293 cells were plated in 10 cm dishes and transfected in biological triplicate at 70% confluency using 10 µg of either mChy-NLS or mChy-ICP4 per plate. 24h after transfection, cells were trypsinized, washed with PBS, and resuspended in sorting buffer (1X PBS, 1mM EDTA, 25 mM HEPES (pH 7.0), 1% FBS) at a density of 5x10^6^ cells/mL. mChy-positive cells were sorted using a BD FACSAria Fusion Cell Sorter, collecting exactly 500,000 cells per transfection. After collection, cells were spun down and resuspended in TRIzol Reagent (ThermoFisher). Total RNA was extracted, DNaseI treated (NEB), and quantified using the Qubit RNA HS Assay Kit (ThermoFisher). One-step qRT-PCR was performed using the Luna Universal One-Step RT-qPCR Kit (NEB) and 5 ng RNA per reaction in technical duplicate with a StepOnePlus Real-Time PCR system (Applied Biosystems) with SYBR Green detection. The expression levels of target mRNAs were first normalized to 18S rRNA then compared across conditions using the ΔΔC_T_ method. Primer sequences are in Supplemental Figure 7.

### Accession numbers

Sequencing data were deposited in the NCBI Gene Expression Omnibus and are accessible through GEO Series accession numbers GSE171600. To analyze the ICP4 ChIP-seq data (Figure 3c) raw reads were downloaded from SRA (accession nos. SRX6422223 and SRX6422222) and mapped to the hg19 assembly as described above.

## Abbreviations

Pol II: RNA polymerase II
HSV-1: Herpes simplex virus type 1
ICP-: infected cell polypeptide
TFII-: transcription factor of Pol II
NELF: negative elongation factor
DSIF: DRB sensitivity inducing factor
PTEF-b: positive transcription elongation factor
CTD: Cterminal domain
ChIP-seq: Chromatin immunoprecipitation followed by sequencing
WT: wildtype
RPK: mapped sequence reads per kilobase
TSS: transcription start site
GO: gene ontology
mChy-: mCherry tagged polypeptide
AP-1: activator protein 1 transcription factor

## Acknowledgements

This work was supported by grants from the National Institutes of Health (R01 GM068414; T.R. was partially supported by the training grants T32 GM008759 and F31 GM125366). We thank Theresa Nahreini and the Flow Cytometry Shared Core Facility (supported by NIH S10ODO21601). We thank the N. DeLuca lab for the n12 virus and complementing cell line, and the L. Schang lab for the mChy-ICP4 plasmid.

