## Supplementary Materials for "The herpes simplex virus 1 protein ICP4 acts as both an activator and repressor of host genome transcription during infection"

Supplemental Figures 1-7

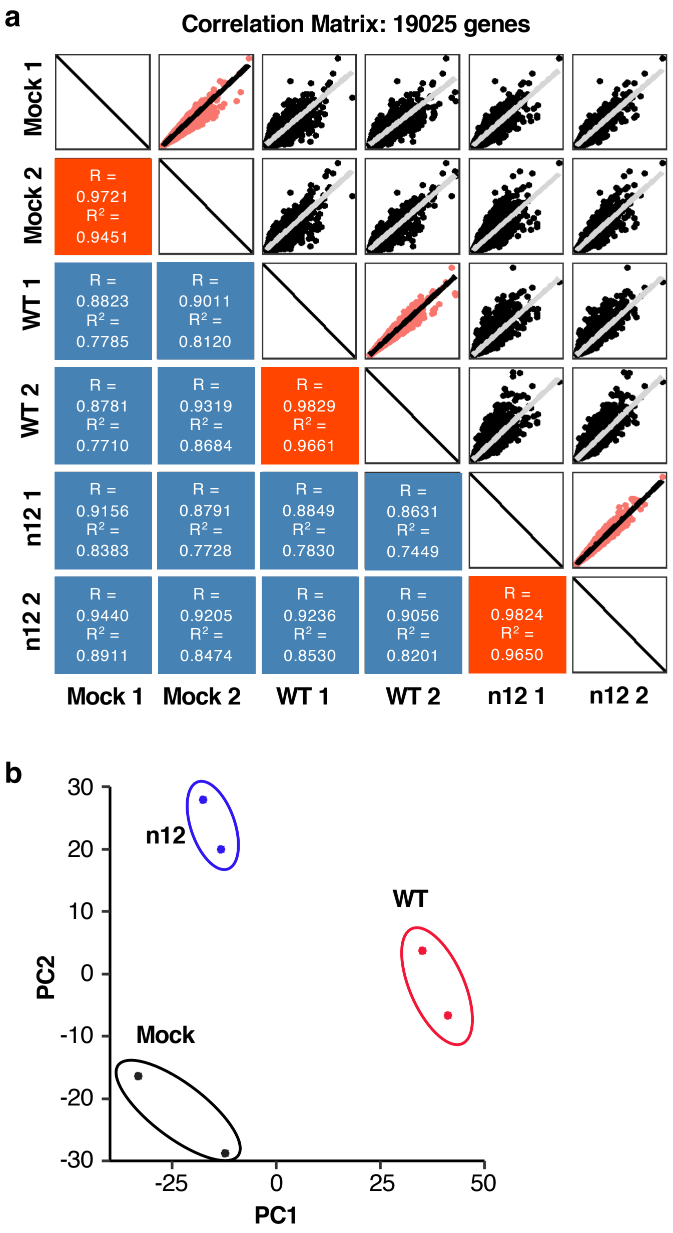

**Supplementary Figure 1.** The ChIP-seq data for biological replicates from each condition cluster with each other, but not with different conditions. **(A)** Shown is a correlation matrix comparing the Pol II ChIP-seq read coverage across 19,025 individual mRNA isoforms from each replicate for each condition. Pearson values (R) and coefficients of determination are shown on the bottom left half and the corresponding scatter plots on the top right half. Biological replicates are highlighted in red. **(B)** Different infection conditions provide distinct Pol II ChIP-seq data. PCA plot showing the relationship of individual replicates to each other. Replicate infection conditions cluster together and are distinct from other infection conditions. PC1 and PC2 explain 59.2% and 29.6% of the variance, respectively.

| ID | Term | Count | % | P-adj value |
|---|---|---|---|---|
| **WT Activated Genes, BP_Direct** | | | | |
| GO:0001701 | *In utero* embryonic development | 19 | 4.36 | 6.8E-04 |
| GO:0000122 | *Negative regulation of transcription from RNA polymerase II promoter | 41 | 9.40 | 6.8E-04 |
| GO:0045944 | *Positive regulation of transcription from RNA polymerase II promoter | 49 | 11.24 | 1.3E-03 |
| GO:0006366 | *Transcription from RNA polymerase II promoter | 31 | 7.11 | 3.4E-03 |
| GO:0016477 | Cell migration | 16 | 3.67 | 7.4E-03 |
| GO:0051591 | Response to cAMP | 8 | 1.83 | 3.8E-02 |
| **n12 Activated Genes, BP_Direct** | | | | |
| GO:0045944 | *Positive regulation of transcription from RNA polymerase II promoter | 48 | 12.53 | 8.04E-05 |
| GO:0006366 | *Transcription from RNA polymerase II promoter | 32 | 8.36 | 8.04E-05 |
| GO:0035914 | Skeletal muscle cell differentiation | 9 | 2.35 | 4.35E-03 |
| GO:0000122 | *Negative regulation of transcription from RNA polymerase II promoter | 34 | 8.88 | 6.99E-03 |
| GO:0007399 | Nervous system development | 18 | 4.70 | 3.87E-02 |
| **WT Repressed Genes, BP_All** | | | | |
| GO:0006807 | Nitrogen compound metabolic process | 258 | 43.95 | 4.73E-03 |
| GO:0008152 | Metabolic process | 373 | 63.54 | 1.04E-02 |
| GO:0034641 | Cellular nitrogen compound metabolic process | 240 | 40.89 | 1.09E-02 |
| GO:0044237 | Cellular metabolic process | 347 | 59.11 | 1.09E-02 |
| GO:1901360 | Organic cyclic compound metabolic process | 228 | 38.84 | 1.09E-02 |
| GO:0071704 | Organic substance metabolic process | 360 | 61.33 | 1.30E-02 |
| GO:0006259 | DNA metabolic process | 53 | 9.03 | 2.75E-02 |
| GO:0006725 | Cellular aromatic compound metabolic process | 218 | 37.14 | 3.79E-02 |

**Supplementary Figure 2.** Genes activated by infection are involved in transcriptional regulation while repressed genes are involved in metabolism. Summary of statistically enriched GO terms for genes differentially expressed upon WT and n12 infection. GO analysis was first performed looking at BP_Direct terms (an abbreviated listed of biological pathway terms) with an FDR value < 0.05. No terms were significantly enriched for repressed genes, thus the GO analysis was repeated with the broader list of biological pathway terms, BP_All. n12 repressed genes had no significant GO-terms. * represent GO IDs that are represented in both WT and n12 activated genes.

| Hallmark Gene Set Name | Count | % | P-adj value |
|---|---|---|---|
| **WT Activated Genes** | | | |
| *TNFA_SIGNALING_VIA_NFKB | 39 | 19.50 | 6.68E-35 |
| *P53_PATHWAY | 27 | 13.50 | 7.31E-20 |
| *UV_RESPONSE_UP | 20 | 12.66 | 2.61E-14 |
| *ESTROGEN_RESPONSE_LATE | 20 | 10.00 | 1.87E-12 |
| HYPOXIA | 17 | 8.50 | 1.34E-09 |
| IL2_STAT5_SIGNALING | 15 | 7.54 | 7.11E-08 |
| *ESTROGEN_RESPONSE_EARLY | 14 | 7.00 | 4.84E-07 |
| HEME_METABOLISM | 13 | 6.50 | 2.59E-06 |
| MTORC1_SIGNALING | 13 | 6.50 | 2.59E-06 |
| ANGIOGENESIS | 6 | 16.67 | 1.37E-05 |
| **n12 Activated Genes** | | | |
| *TNFA_SIGNALING_VIA_NFKB | 30 | 15.00 | 8.29E-25 |
| *P53_PATHWAY | 22 | 11.00 | 2.08E-15 |
| *UV_RESPONSE_UP | 18 | 11.39 | 4.91E-13 |
| *ESTROGEN_RESPONSE_LATE | 16 | 8.00 | 2.22E-09 |
| APOPTOSIS | 11 | 6.83 | 6.29E-06 |
| MYOGENESIS | 12 | 6.00 | 6.53E-06 |
| WNT_BETA_CATENIN_SIGNALING | 6 | 14.29 | 2.38E-05 |
| *ESTROGEN_RESPONSE_EARLY | 11 | 5.50 | 3.26E-05 |
| MYC_TARGETS_V2 | 6 | 10.34 | 1.25E-04 |
| INTERFERON_GAMMA_RESPONSE | 10 | 5.00 | 1.58E-04 |

| Hallmark Gene Set Name | Count | % | P-adj value |
|---|---|---|---|
| **WT Repressed Genes** | | | |
| E2F_TARGETS | 17 | 8.50 | 6.61E-07 |
| G2M_CHECKPOINT | 12 | 6.00 | 1.52E-03 |
| CHOLESTEROL_HOMEOSTASIS | 7 | 9.46 | 2.21E-03 |
| GLYCOLYSIS | 11 | 5.50 | 3.26E-03 |
| UNFOLDED_PROTEIN_RESPONSE | 8 | 7.08 | 3.37E-03 |
| MITOTIC_SPINDLE | 10 | 5.03 | 8.19E-03 |
| FATTY_ACID_METABOLISM | 8 | 5.06 | 2.10E-02 |
| MTORC1_SIGNALING | 9 | 4.50 | 2.27E-02 |
| ANDROGEN_RESPONSE | 6 | 6.00 | 2.35E-02 |
| BILE_ACID_METABOLISM | 6 | 5.36 | 3.65E-02 |

**Supplementary Figure 3.** Genes activated by WT and n12 infection are enriched in similar pathways. Shown is a summary of enriched Hallmark gene sets for differentially expressed genes upon WT and n12 infection. The top 10 gene sets with FDR values < 0.05 are listed. No gene sets were enriched for n12 repressed genes. * represent Hallmark gene sets that are represented in both WT and n12 activated genes.

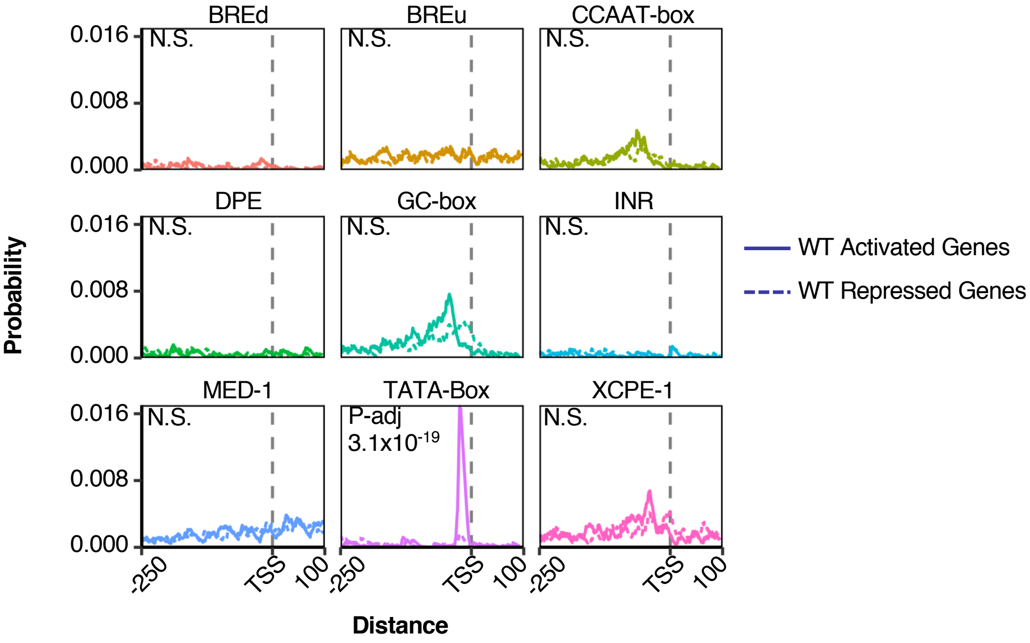

**Supplementary Figure 4.** The TATA-box is enriched in the promoters of HSV-1 activated but not repressed genes. We analyzed for enrichment of nine core promoter elements in the 463 WT activated (solid lines) and 626 WT repressed (dashed lines) genes using CentriMo (-1000 to +100 relative to the TSS). Only the JASPAR Pol II motifs were analyzed.

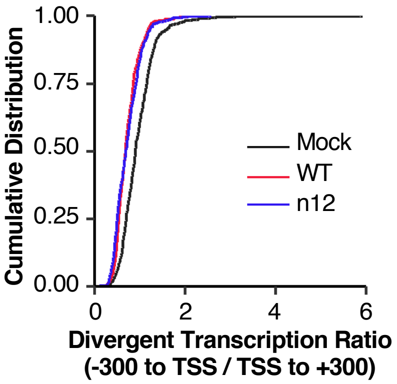

**Supplementary Figure 5.** Divergent transcription is dysregulated upon HSV-1 infection independent of ICP4. Plotted is a cumulative distribution of the $\log_2$ divergent transcription ratio (RPK -300 to TSS/RPK TSS to +300) for the 463 genes activated by WT virus infection, for the mock, WT, and n12 conditions. P-values were determined using a two-sided Kolmogorov-Smirnov test. P-values: mock versus WT, $< 2.2 \times 10^{-16}$; mock versus n12, $< 2.2 \times 10^{-16}$; WT versus n12, 0.075.

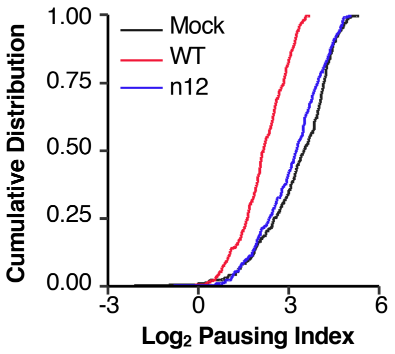

**Supplementary Figure 6.** ICP4 regulates promoter proximal pausing at genes dependent on ICP4 for activation. Plotted is a cumulative distribution of the $\log_2$ pausing indices (gene body RPK/promoter proximal RPK) for the 294 genes dependent on ICP4 for activation upon WT infection, for the mock, WT, and n12 conditions. P-values were determined by using a two-sided Kolmogorov-Smirnov test: mock versus WT, $p < 2.2 \times 10^{-16}$, mock versus n12: $p < 0.0050$; WT versus n12, $p < 2.2 \times 10^{-16}$.

| Name | Sequence |
|---|---|
| EPHA4 GB F | GGAACCCAGCCAGAATAACTGGC |
| EPHA4 GB R | CATGACGCCCGGAAGACTATTGC |
| INCENP GB F | GCAGAAAGTCTCGGAGCAGCC |
| INCENP GB R | GCAGGACGGAGCCGTTCT |
| FADD GB F | GAAAGATTGGAGAAGGCTGGCTCG |
| FADD GB R | CCAGATTCTCAGTGACTCCCGC |
| IGF2 GB F | GGAAGTCGATGCTGGTGCTTCT |
| IGF2 GB R | CAGACGAACTGGAGGGTGTCC |
| EGR3 GB F | GGACATCGGTCTGACCAACGAG |
| EGR3 GB R | GAGTCGAAGGCGAACTTTCCCAAG |
| 18S F | GTAACCCGTTGAACCCCA |
| 18S R | CCATCCAATCGGTAGTAGC |
| VEGFA GB F | CTGCCATCCAATCGAGACCCTG |
| VEGFA GB R | CCCTCGTCATTGCAGCAGC |
| 3xAP-1 QC F | TCGAGCTCGATGGGGATCCGCTAGCTACCCTCTAGAAGCTTGCTGAG |
| 3xAP-1 QC R | CTCAGCAAGCTTCTAGAGGGTAGCTAGCGGATCCCCATCGAGCTCGA |
| EPHA4 PPR F | GGATAGAAGCGGCAGGAGCAG |
| EPHA4 PPR R | CCCTGGAACCTGTGACAGCG |

**Supplementary Figure 7.** Sequences of primers used, written 5’ to 3’. The first group of primers are gene body (GB) primers to analyze mRNA levels by qPCR after sorting. The QuikChange (QC) mutagenesis primers were used to mutate the TATA box in the 3xAP-1 luciferase plasmid. The EPHA4 primers were used to analyze NELF-A, Spt5, and Pol II occupancy in the promoter proximal region downstream of the TSS.
